# Entrainment of circadian rhythms depends on firing rates and neuropeptide release of VIP SCN neurons

**DOI:** 10.1101/322339

**Authors:** Cristina Mazuski, John H. Abel, Samantha P. Chen, Tracey O. Hermanstyne, Jeff R. Jones, Tatiana Simon, Francis J. Doyle, Erik D. Herzog

**Affiliations:** Department of Biology, Washington University in St. Louis, St. Louis, MO 63130; Department of Systems Biology, Harvard Medical School, Boston, MA 02115; Harvard John A. Paulson School of Engineering and Applied Sciences, Harvard University, Cambridge, MA 02138

**Author notes:** Corresponding author/Lead Contact.

**Keywords:** suprachiasmatic nucleus, Period gene, optogenetics, channelrhodopsin-2, multielectrode array, electrophysiology, daily oscillation

## Abstract

The mammalian suprachiasmatic nucleus (SCN) functions as a master circadian pacemaker, integrating environmental input to align physiological and behavioral rhythms to local time cues. Approximately 10% of SCN neurons express vasoactive intestinal polypeptide (VIP); however, it is unknown how firing activity of VIP neurons releases VIP to entrain circadian rhythms. To identify physiologically relevant firing patterns, we optically tagged VIP neurons and characterized spontaneous firing over three days. VIP neurons had circadian rhythms in firing rate and exhibited two classes of instantaneous firing activity. We next tested whether physiologically relevant firing affected circadian rhythms through VIP release. We found that VIP neuron stimulation with high, but not low, frequencies shifted gene expression rhythms *in vitro* through VIP signaling. *In vivo*, high frequency VIP neuron activation rapidly entrained circadian locomotor rhythms. Thus, increases in VIP neuronal firing frequency release VIP and entrain molecular and behavioral circadian rhythms.

**Highlights:**

- Mazuski et al. identified three classes of circadian SCN neurons based on their distinct firing patterns consistent over multiple days
- There are two distinct classes (tonic and irregular firing) of VIP SCN neurons.
- Stimulation of VIP SCN neurons at physiologically relevant frequencies phase shifts whole-SCN circadian rhythms in gene expression through VIP release. These effects are blocked with VIP antagonists.
- Firing of VIP SCN neurons entrains circadian rhythms in locomotor behavior in a frequency and time-of-day dependent manner.

## Introduction

Circadian rhythms are a ubiquitous adaptation to the 24-h daily cycle on Earth. Found in organisms as disparate as *Arabidopsis, Drosophila, Synechococcus*, and humans, these daily oscillations occur at molecular, cellular and systems levels, ultimately entraining an organism to environmental cycles(Dunlap, 1999). Dysregulation in the circadian system, also known as chronodisruption, is associated with a variety of physiological maladies in humans including diabetes, cancer, insomnia, and affective disorders(Van Someren, 2000, Musiek et al., 2013, McClung, 2011). Understanding how cellular circadian oscillators integrate environmental signals, communicate with each other, and entrain to their environment is key to preventing and treating chronodisruption.

The suprachiasmatic nucleus (SCN) is a master circadian pacemaker, which detects local light input through retinal release of glutamate and PACAP(Eastman et al., 1984, Ding et al., 1997, Harrington et al., 1999, Hannibal et al., 2000). Located in the ventral hypothalamus, the approximately 20,000 GABAergic neurons comprising the SCN are unique in that many express self-sustained and spontaneously synchronizing circadian rhythms in firing activity and gene expression(Cassone et al., 1993, Reppert and Weaver, 2002, Welsh et al., 1995). These rhythms are maintained through a near-24 h transcription-translation feedback loop of core clock genes including *Bmal1, Clock, Period1 and 2 (Per1 and 2)*, and *Cryptochrome1 and* 2(Ko and Takahashi, 2006, Okamura et al., 2002, Takahashi, 2017). Recently, researchers have shown that electrical stimulation of all SCN neurons was sufficient to phase shift and entrain circadian rhythms in gene expression and behavior(Jones et al., 2015). However, it is poorly understood how firing activity and neurotransmission from specific cell types within the SCN contribute to entrainment.

Although uniformly GABAergic (Moore and Speh, 1993), SCN neurons vary significantly in their neuropeptide content (Abrahamson and Moore, 2001). One anatomically and functionally distinct class of neurons expresses vasoactive intestinal polypeptide (VIP). Though neurons expressing VIP only make up approximately 10% of SCN neurons (Abrahamson and Moore, 2001), VIP neuron projections densely innervate the SCN and the majority of SCN neurons express its receptor VPAC2 (An et al., 2012). Functionally, genetic loss of VIP or VPAC2R weakens synchrony among SCN neurons and dramatically reduces circadian rhythms in the SCN and in behavior (Aton et al., 2005, Maywood et al., 2006, Brown et al., 2007). Additionally, exogenous application of VIP increases GABAergic neurotransmission (Reed et al., 2002), induces clock gene expression (Nielsen et al., 2002), and phase shifts daily rhythms in the SCN and locomotor activity (An et al., 2011, Piggins et al., 1995). With the development of Cre-lox technology allowing cell-type specific manipulation of SCN neurons, researchers found that VIP neurons have sparse GABAergic monosynaptic connections to approximately 50% of SCN neurons (Fan et al., 2015).

Despite the important role VIP plays in circadian function, we do not know how VIP is released from SCN neurons other than it is stored within dense core vesicles (Castel et al., 1996). The release of neuropeptides has been studied primarily in large neuromuscular synapses and hippocampal neurons where firing rates in excess of 50Hz have been associated with release (Xia et al., 2009, Pecot-Dechavassine and Brouard, 1997). In contrast, SCN neurons fire at sustained rates that rarely exceed 15Hz (Welsh et al., 1995). Therefore it remains unclear what role, if any, firing activity in VIP neurons plays in neuropeptide release and circadian entrainment.

To address whether increases in VIP neuronal firing frequency mediate circadian entrainment, we took a multi-step approach. We first characterized the spontaneous, daily firing frequencies and patterns of VIP neurons. We then stimulated VIP neurons comparing the effects of different firing frequencies on daily rhythms *in vivo* and *in vitro*.

## Results

### Optical tagging reveals stable firing patterns of VIP SCN neurons over multiple days

Single SCN neurons exhibit changes in spontaneous firing activity across the circadian day that can vary dramatically in frequency and firing pattern. It is not known whether these electrophysiological characteristics relate to cell-intrinsic properties like neuropeptide release. Therefore, we designed our experiment to characterize firing activity of neuropeptidergic VIP SCN neurons by recording from single neurons across multiple days.

To do so, we optically tagged VIP neurons within SCN cultures plated on multielectrode arrays (MEAs). Specifically, following multiday spontaneous activity recordings, we activated channelrhodpsin-2 (ChR2) expressed solely within VIP neurons (VIP-IRES-Cre crossed with floxed-ChR2, see methods) using blue light pulses (Figure 1a). We identified 9.1 ± 2.4% (mean ± SEM) of recorded neurons as VIP neurons because they had an interspike interval histogram (ISIH) during the 1 h stimulation that matched the stimulation pattern (Figure 1b). We then cross-correlated firing between each pair of activated neurons and found they reliably fired synchronously in response to stimulation. These VIP neurons contrasted with neurons which either did not change or decreased their firing approximately 10-20 ms following stimulation, consistent with inhibitory neurotransmission from VIP SCN neurons (Figure 1c).

**Figure 1.**
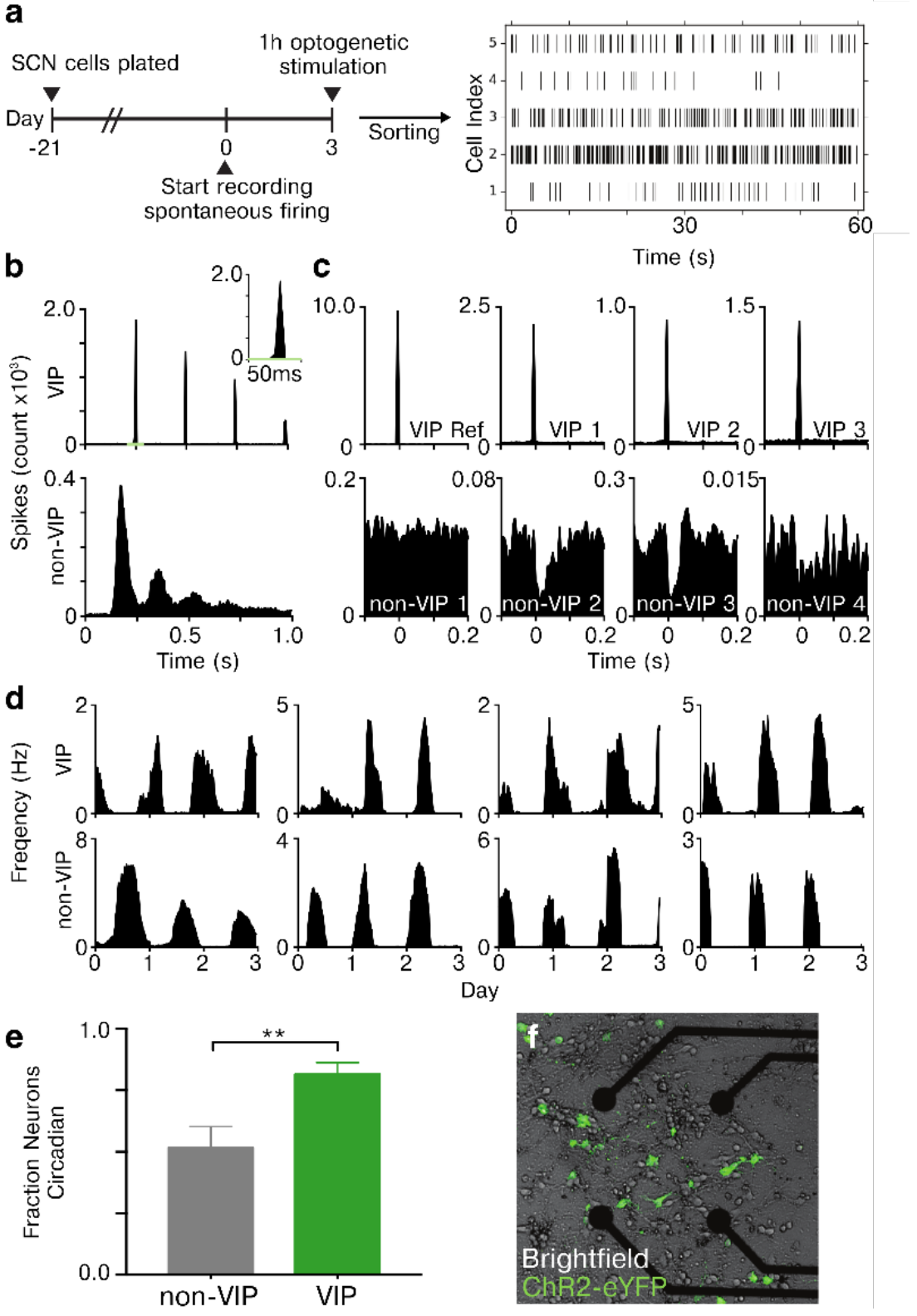
Characterizing multiday spontaneous firing activity of identified VIP SCN neurons within a multielectrode array culture. **a)** SCN neurons were categorized as VIP-positive (VIP) or negative (non-VIP) by optically tagging VIP neurons using optogenetic stimulation after 3 days of spontaneous activity recording. Multiday firing was sorted from 582 SCN neurons identified on 10 multielectrode arrays plated and cultured for 3 weeks from VIPChR2 mice. Raster plots of five representative SCN neurons show how their spike times over one minute differed in mean rate and pattern. **b)** The inter-spike interval histograms during optogenetic stimulation illustrate how a representative VIP (top) neuron fired at the stimulation frequency (4 Hz) with a precision of <10ms (top right inset) and a non-VIP neuron (bottom) fired in a ChR2-independent pattern. **c)** To further characterize the evoked firing of VIP neurons, we cross-correlated spike times between concurrently recorded SCN neurons during optogenetic stimulation. A VIP reference neuron (top left panel) fired synchronously with 3 other representative VIP (top panels), but not 4 representative non-VIP neurons (bottom panels). Note that some non-VIP neurons (#2 and #3) decreased their firing following stimulation of VIP neurons, indicative of postsynaptic inhibition. **d)** Four representative VIP (top) and non-VIP (bottom panels) SCN neurons showing circadian firing patterns over the three days of recording. **e)** A greater fraction of VIP neurons were circadian (81.9 ± 6.2%) compared to non-VIP neurons (51.7 ± 8.5%, Chi-squared test **p < 0.00001). **f)** eYFP fluorescence (green) reveals the subset of SCN neurons expressing ChR2 near four of the 60 electrodes. For further verification of optical tagging of VIP neurons see Figure S1.

We next characterized the spontaneous activity of VIP and non-VIP neurons. Overall, we discriminated multiday electrical activity from 42 VIP neurons and 540 non-VIP neurons across 10 MEA preparations. Two prior reports both measured firing for 1-30 min at two times of day (Hermanstyne et al., 2016, Fan et al., 2015), but differed on whether VIP SCN neurons have circadian firing rhythms. By following individual VIP neurons across multiple days, we found that a majority had circadian firing activity (81.9 ± 6.2%, Figure 1d-1e). After optical tagging, we verified that a subset of SCN neurons expressed ChR2-eYFP (figure 1f), consistent with VIP-neuron specific recombination (Hermanstyne et al., 2016, Fan et al., 2015, Brancaccio et al., 2013). Parallel whole-cell patch clamp recordings in slices and MEA recordings from dissociated cultures revealed that only ChR2-positive SCN neurons fired within 5 ms after light flashes (Figure S1). In conclusion, optical tagging reliably identified the multiday, circadian firing of VIP SCN neurons.

### Evidence for two types of VIP SCN neurons

Most electrophysiological SCN studies have focused on circadian changes in SCN neuronal firing rate (the number of spikes in bins ranging from 10s to 10min; Figure 1d). Because neurotransmitter release occurs on millisecond timescales, we instead elected to analyze instantaneous spike timing and frequency of circadian SCN neurons (36 VIP and 302 non-VIP).

Based on the spike timing of spontaneous activity during an individual neuron’s peak daily firing, we observed three distinct classes of spike timing within SCN neurons: tonic, irregular, and bursting (Figure S2) similar to those previously described (Pennartz et al., 1998). Unsupervised, hierarchical clustering based on the ISIH and dominant instantaneous firing rate (DIFR) revealed that VIP neurons could be classified as having either tonic or irregular spike timing (Figure 2a) whereas non-VIP neurons were tonic, irregular or exhibited short bursts of firing greater than 50Hz (Figures 2b-2c). The DIFR did not differ between tonic and irregular VIP or non-VIP neurons, but distinguished the bursting non-VIP neurons of the SCN (Figure 2d). Strikingly, individual SCN neurons did not change their stereotyped spike timing (tonic, irregular, or bursting) with time of day, or across multiple days, including during periods of rapidly increasing or decreasing firing activity (Figure 2e). We found two comparable classes when analyzing intracellular recordings from VIP neurons in adult SCN slices (Figures 2f-2h, data originally collected in Hermanstyne et al. 2016). These results indicate that VIP SCN neurons are circadian and, despite dynamic fluctuations in their mean firing rate, can be classified as either tonic or irregular spiking independent of time of day over multiple days.

**Figure 2.**
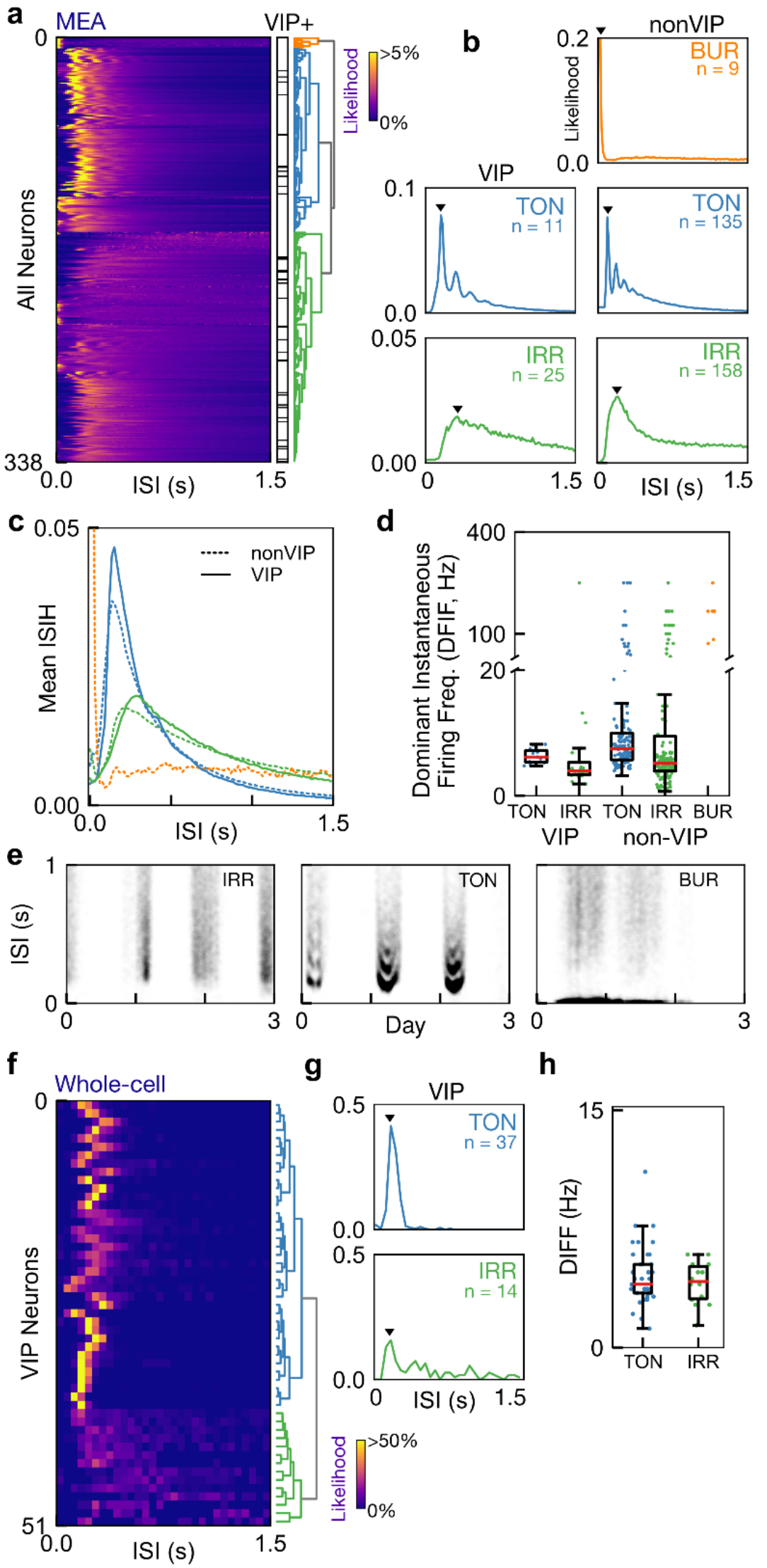
VIP SCN neurons exhibit tonic or irregular circadian firing patterns. **a)** Unsupervised hierarchical clustering sorts circadian SCN neuron recordings from multielectrode arrays into one of three different daytime firing patterns: Tonic (TON), irregular (IRR) or bursting (BUR). Identified VIP neurons (black lines) formed a heterogeneous class that exhibited either tonic or irregular firing. See also Figure S2. **B)** Representative interspike interval histograms (ISIH) for each identified class of neurons. The dominant instantaneous firing rate (DIFR, black triangle) measures the most common firing interval for an individual neuron. (n = number of neurons recorded within each class) **C)** We found no differences within each firing class (colored lines as in B) when comparing the average ISIH of VIP and non-VIP neurons. **d)** Bursting cells had higher DIFR than other cell classes (median (interquartile range) in Hz of TON VIP 6.2 (1.9), IRR VIP 3.9 (2.0), TON Non-VIP 7.5 (4.3), IRR Non-VIP 5.1 (5.5), and BUR Non-VIP 166.6 (83.3)). **e)** Short-term firing patterns were stable over three days shown by three representative SCN neurons. Note, for example, the daily appearance of multiple bands during tonic firing, corresponding to harmonics resulting from skipped spikes. **f)** Daytime firing VIP neurons recorded from SCN slices using whole-cell patch clamp (data previously published in Hermanstyne et al. 2016) exhibited either tonic or irregular firing similar to MEA recordings based on **g)** ISIH and **h)** DIFR distributions (all values median (interquartile range): tonic VIP 4.0 (4.2) Hz and irregular VIP 4.2 (2.0) Hz).

### High frequency firing of VIP neurons phase shifts circadian gene expression rhythms *in vitro* through VIP release

To test how VIP neuronal firing influences circadian entrainment, we optogenetically stimulated VIP neurons with one of two firing frequencies that represent their high and low frequency ranges: high instantaneous frequency (HIF, doublets of 20 Hz at 2 Hz) or low instantaneous frequency (LIF, tonic 4 Hz; Figures 3a-b). These activation patterns evoked the same number of spikes per second (4 Hz), but differed in their spike timing for the duration of the 1 h of stimulation (Figure S3).

We found that HIF, but not LIF, shifted circadian gene expression in SCN slices. Briefly, we cultured SCN explants expressing ChR2 in VIP neurons and *Period2::Luciferase* (*PER2::LUC*, a knock-in fusion that reports PER2 protein abundance) or littermate controls lacking ChR2 expression. Following baseline bioluminescence recording of PER2 levels from the entire SCN, we stimulated all SCN for 1 h near the peak of PER2 expression (circadian time, CT 9-12) with HIF or LIF patterns. We chose to stimulate at this time because it has been reported as the time when exogenous VIP application evoked large delays in PER2 rhythms (An et al., 2011). HIF stimulation significantly delayed the daily rhythms of SCN PER2 expression by over 1.5 h, whereas LIF stimulation did not (Figures 3c-d). Additionally, three consecutive days of stimulation yielded similar results (Figure S4). Thus, firing of VIP neurons can phase delay circadian gene expression but only if stimulated at sufficiently high frequencies.

**Figure 3.**
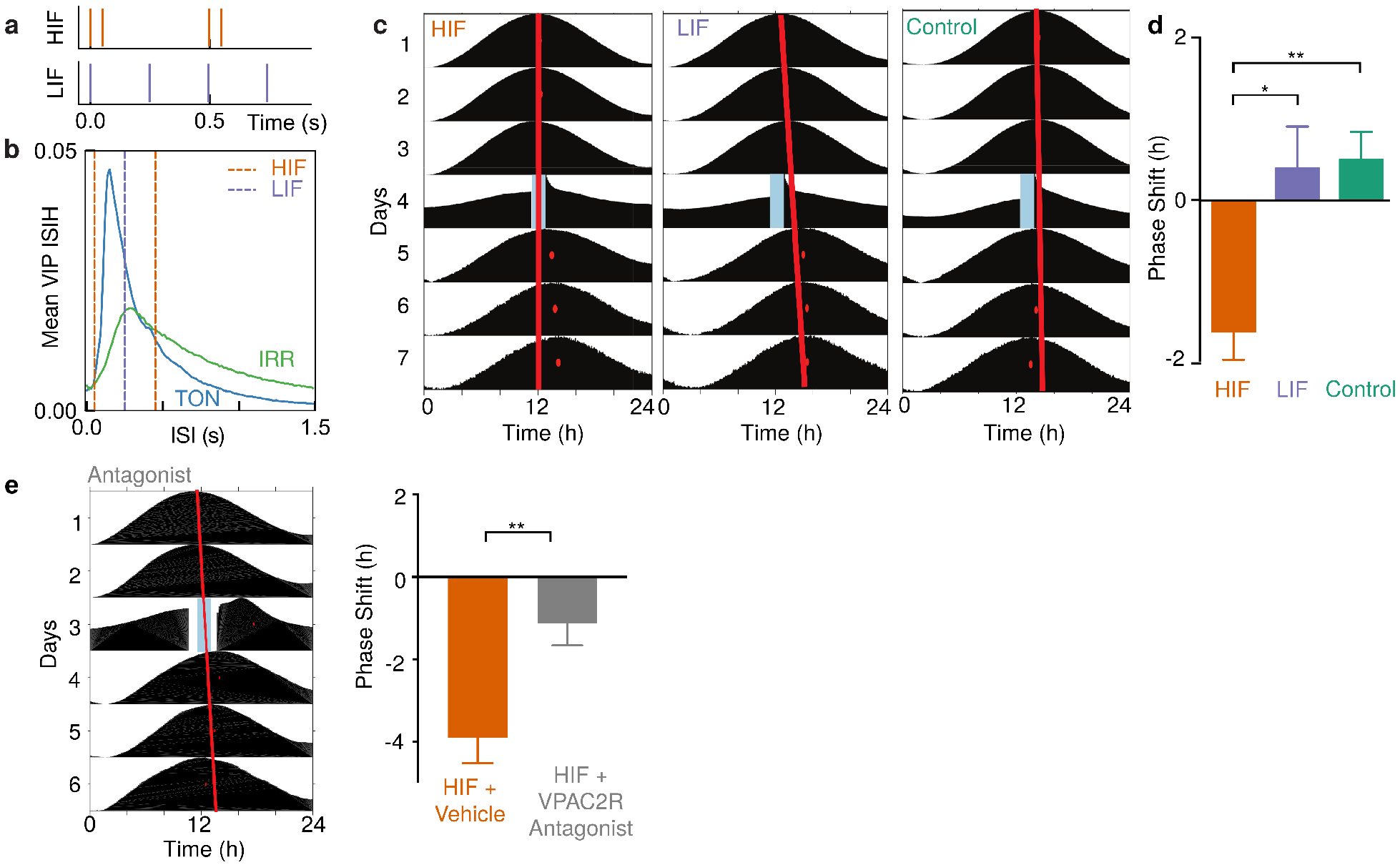
High frequency optogenetic stimulation of VIP SCN neurons phase delays circadian rhythms in PER2 expression. **a)** Based on the spontaneous firing observed in VIP neurons, we used either high instantaneous frequency (HIF) and low instantaneous frequency (LIF) optogenetic stimuli to evoke the same number of action potentials with substantially different interspike intervals. See also Figure S3. b) HIF and LIF optogenetic stimulation frequencies (dashed lines) superimposed on the average ISIH of VIP SCN neurons from Figure 2C. The selected stimulation frequencies represent physiological ranges of VIP firing frequencies irrespective of firing class. **c)** Representative PER2::LUC actograms from three SCN slices. In the first two traces, ChR2 expressed in VIP neurons was activated for 1 h (blue bar) on the fourth day of recording with high (HIF) or low (LIF) frequencies. Control SCN received either HIF or LIF stimulation, but lacked ChR2. Note the large delay in the time of the daily peak of PER2 (red dots) on the days after HIF stimulation relative to the extrapolated unperturbed phase (red line). **d)** HIF stably delayed PER2 rhythms compared to either LIF or control (−1.6 ± 0.3 h, HIF, 0.4 ± 0.5, LIF, 0.5 ± 0.3, control; *p < 0.05 and **p < 0.01, one-way ANOVA with Tukey’s posthoc test, n= 9, 10 and 32 SCN slices, respectively). See also Figure S4. **e)** VPAC2 receptor antagonist treatment during HIF stimulation reliably reduced the resulting phase shift, shown by a representative actogram (left) and the group summary. (mean ± SEM: −3.9 ± 0.62, HIF + Vehicle, −1.1 ± 0.54, HIF + VPAC2R Antagonist; **p < 0.01, unpaired Student’s t-test, n = 7, 7 SCN slices, respectively). See also Figure S5.

To test whether VIP neuronal stimulation causes VIP release, we engineered sensor cells to measure VIP following HIF or LIF optogenetic stimulation (Figure S5). We found that acute HIF activation more potently increased VIP release than acute LIF stimulation. Critically, we found that 10 uM VPAC2R antagonist applied during HIF stimulation reduced the resulting phase shift in SCN explants expressing ChR2 in VIP neurons (Figures 3e-f). Thus, our data suggest that HIF firing of VIP neurons releases VIP which phase delays circadian gene expression.

### Activation of VIP neurons induces cFOS expression throughout the SCN *in vivo*

Although VIP neuronal cell bodies localize to the ventral SCN, VPAC2 receptor is expressed in nearly all SCN neurons (An et al., 2012, King et al., 2003). Consistent with these anatomical data, we found that activation of VIP neurons *in vivo* increases cFOS protein throughout the SCN. Briefly, freely moving mice expressing ChR2 in VIP neurons (VIPChR2) and littermate controls received blue laser light via an implanted fiber optic cannula aimed at the SCN. After 1 h of 15Hz stimulation at CT 13, 88.9 ± 4.0% (mean ± SEM, n = 4 mice; Figure 4a) of VIP neurons expressed cFOS, a marker of neuronal activation. Furthermore, cFOS expression increased roughly 5-fold in the ventral SCN and almost 4-fold in the dorsal SCN (Figure 4b) compared to controls. We conclude that high frequency *in vivo* stimulation of VIP neurons increases activity both within VIP neurons and activates the whole SCN.

**Figure 4.**
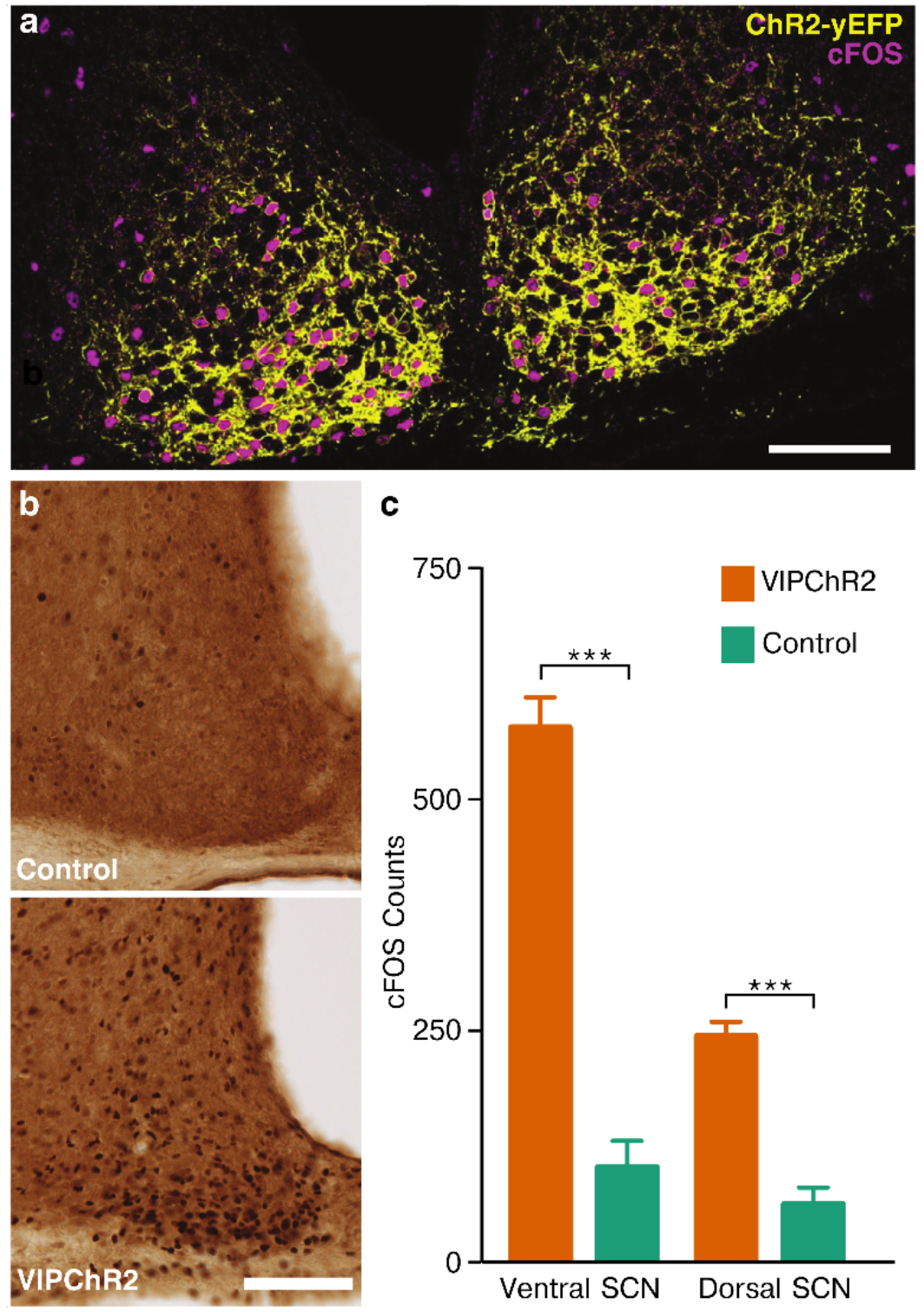
Activation of SCN VIP neurons *in vivo* induces cFOS expression throughout the SCN. **a)** Representative image of the bilateral SCN showing cFOS induction (magenta) within VIPChR2 neurons (yellow) after 1 h of 15 Hz stimulation *in vivo* at CT13 (scale bar = 100um). **b)** The number of SCN cells expressing cFOS protein was higher in VIPChR2 (bottom) than control (top panel) mice after 1 h of 15 Hz stimulation at CT13. *In vivo* optogenetic stimulation of VIP neurons increased cFOS expression throughout the SCN (mean±SEM: 577.8 ± 33.0, Ventral SCN VIPChR2, 103.0 ± 56.9, Ventral SCN Control, 244.5 ± 15.6 Dorsal SCN VIPChR2, 63.0 ± 17.9 Dorsal SCN Control; unpaired Student’s t-test ***p<0.001, n= 4 mice).

### Activation of VIP neurons entrains locomotor activity *in vivo*

Next, we found that firing of VIP neurons underlies their role in circadian entrainment. Briefly, we monitored wheel-running activity before, during and after stimulation of VIP SCN neurons *in vivo* from enucleated mice (to block ambient light input) implanted with a fiber optic directed at the SCN. Before stimulation, all mice showed similar free-running circadian rhythms in locomotion (in hours, 23.4 ± 0.1, VIPChR2, 23.4 ± 0.2, Control, mean ± SEM, n = 7, 4). Mice then received daily HIF stimulation for 1 h for up to 30 days and only VIP-ChR2 mice entrained to the stimulation (Figure 5a). We calculated a phase response curve (Figure 5b) and found that stimulation of VIP neurons between CT 8-18 led to phase delays and entrainment. 10 Hz activation of VIP SCN neurons for 1 h at the same time each day similarly entrained circadian locomotor behavior (Figure S6). Additionally, we observed that stimulation of VIP neurons acutely suppressed wheel running (Figure 5c-d) similar to light-induced masking of locomotion.

**Figure 5.**
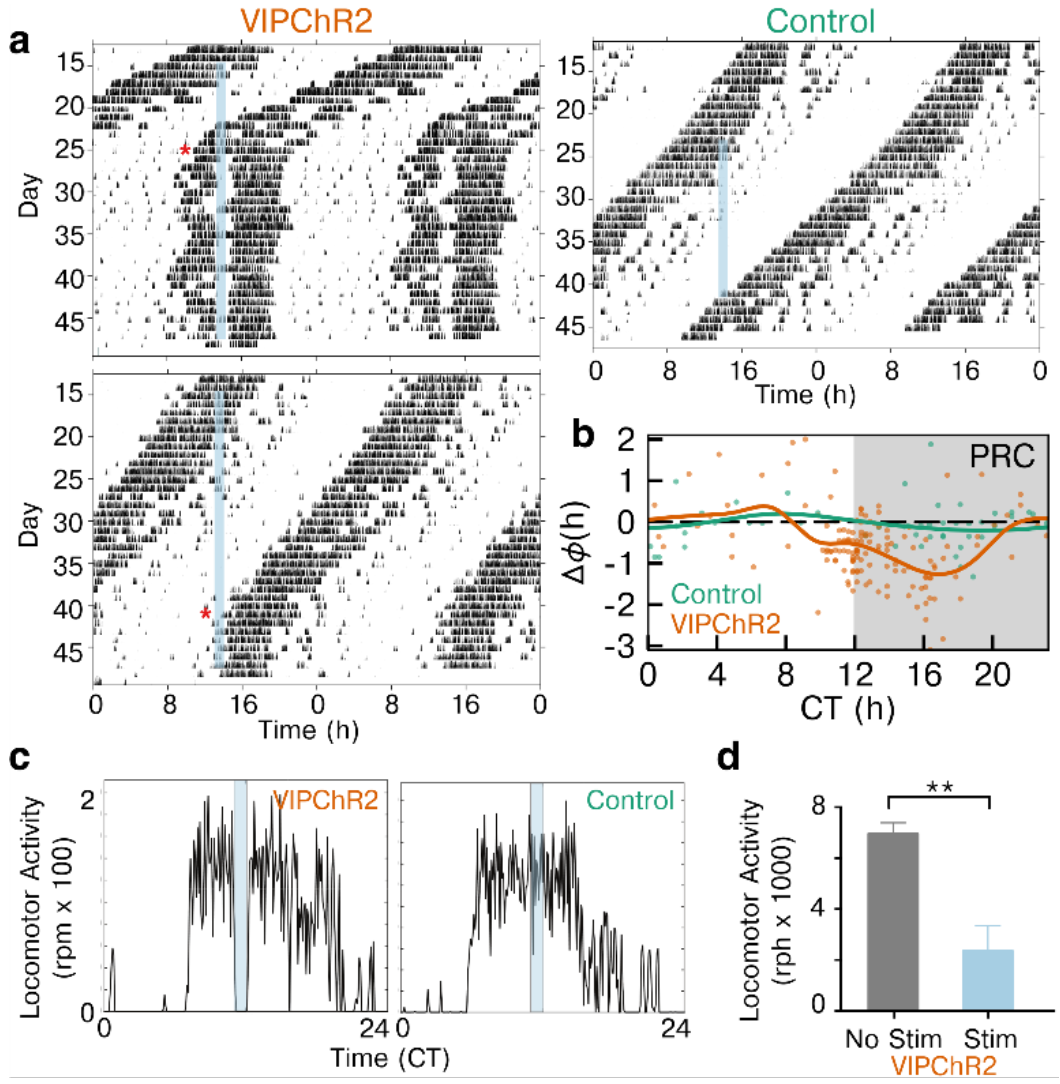
Stimulation of VIP neurons in vivo entrains locomotor activity. **a)** Daily locomotor activity of two representative mice entrained to HIF stimulation of SCN VIP neurons (blue bar) compared to a control mouse lacking ChR2. Actograms show wheel revolutions per 6 min (black bars) recorded from enucleated mice for almost 50 days. Note that the two VIPChR2 mice, with slightly different periods, reached stable entrainment (*) only when the stimulation occurred around early subjective night. **b)** Average phase response curves following stimulation of VIPChR2 (orange, n = 7) and control mice (green, n = 4) show the change in phase (dots) on the day after stimulation at different circadian times. Note that activation of VIP neurons entrained daily locomotor rhythms primarily through phase delays when delivered during the late subjective day and early subjective night. **c)** Representative activity profiles of 2 mice show that HIF stimulation of VIP neurons acutely reduced running wheel activity in VIPChR2 (left), but not control (right), mice. **d)** Wheel revolutions during optogenetic stimulation (CT 12-18, 1 h of HIF) decreased nearly fourfold compared to baseline in VIPChR2 mice (2355.0 ± 981.1 during stimulation vs. 6958.0 ± 418.9 with no stimulation, n = 6 mice, paired Student’s t-test **p< 0.01).

Finally, we found that firing frequency of VIP SCN neurons plays a role in the rate of circadian entrainment. Mice were randomly assigned to receive either 1h of daily HIF or LIF stimulation for 4-10 consecutive days. We found that either stimulation frequency entrained locomotor activity (Figure 6c), however HIF stimulation entrained the mice more rapidly (Figure 6a-b). We conclude that firing of VIP neurons results in circadian entrainment and the frequency of firing determines the speed of entrainment.

**Figure 6.**
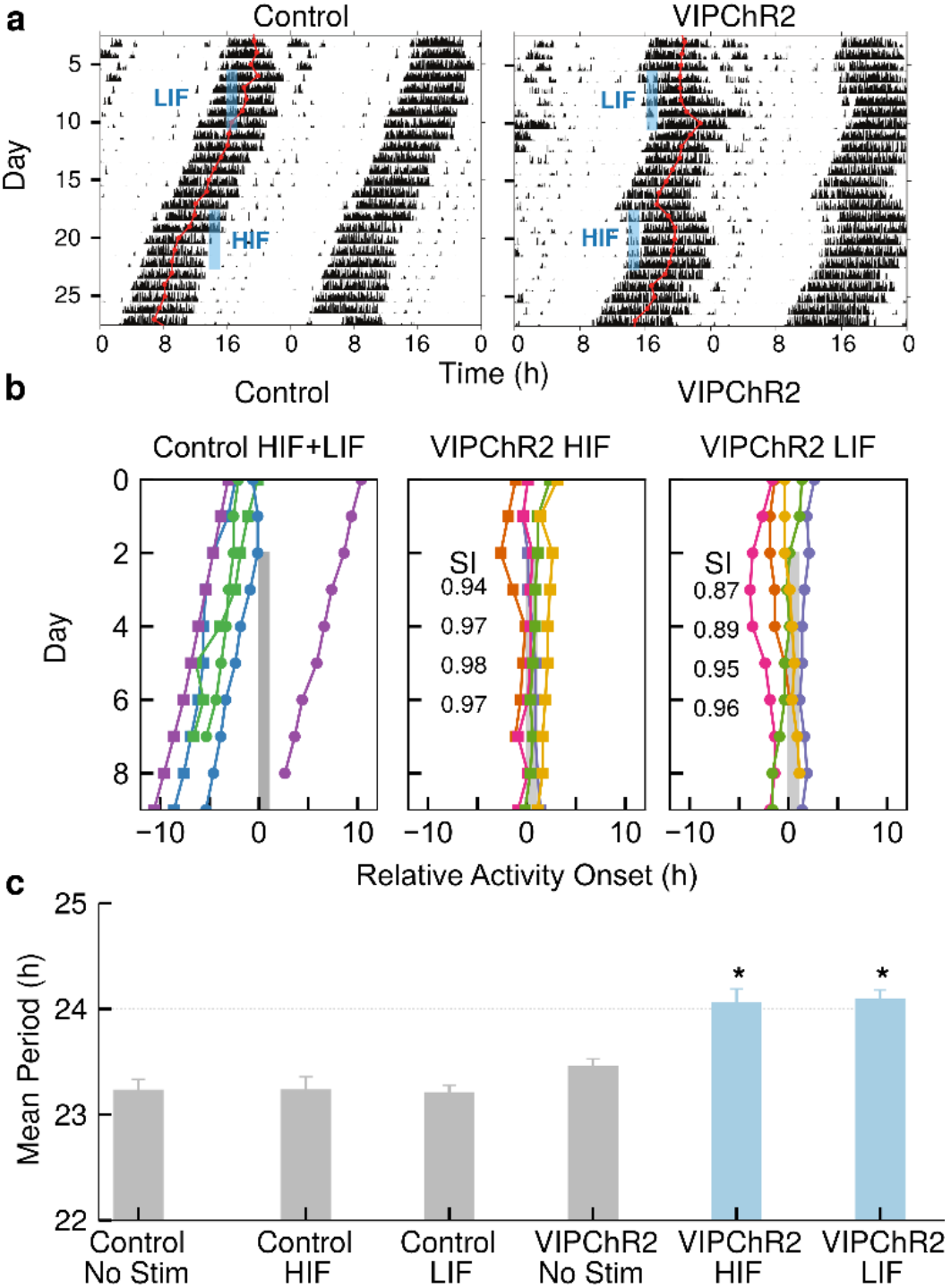
Instantaneous firing pattern affects locomotor rhythm entrainment. **a)** Representative actograms show how daily optogenetic stimulation with HIF or LIF (blue bar) differentially entrained VIPChR2, but not control, mice. The daily acrophase (red circles and lines) of control mice in constant darkness free-ran through the days of stimulation. In contrast, daily HIF stimulation produced a large delay and rapidly entrained locomotor rhythms. Daily LIF stimulation, while having smaller effects on phase, also entrained. **b)** Daily stimulation entrained daily activity onsets to within 2 h of the stimulation in both HIF and LIF but not in control mice. HIF stimulation immediately resulted in tight clustering of activity onsets, as shown by the higher synchrony index (SI), while LIF stimulation gradually entrained mice to the stimulation, as seen by the gradually increasing SI. Each colored line in HIF and LIF conditions indicates data from one mouse. **c)** Daily HIF or LIF stimulation synchronized locomotor rhythms to 24 h in VIPChR2, but not control, mice (one-way ANOVA with posthoc Tukey HSD *p < 0.05). Note that VIPChR2 mice displayed a period identical to controls while stimulation was off. See also Figure S6.

## Discussion

### VIP SCN neurons are electrophysiologically diverse

We consistently found that VIP SCN neurons fired with either tonic or irregular patterns in extracellular, multiday and intracellular, acute-slice recordings. We hypothesize that these firing classes relate to two functional classes of VIP SCN neurons, in agreement with two prior anatomical studies that postulated two groups of VIP SCN neurons based on their differential, light-evoked, *Period1* gene expression (Kawamoto et al., 2003) and developmental expression of *Vip* mRNA (Ban et al., 1997). Intriguingly, these may represent VIP neurons that either project within or outside the SCN (Abrahamson and Moore, 2001). Similar subgroups of brainstem serotonergic neurons have been associated, for example, with regulating distinct behaviors including respiration and aggression through distinct brain target areas (Niederkofler et al., 2016). Future studies should elucidate whether the physiological classes of VIP and non-VIP neurons map to specific anatomical groups within the SCN and targets outside the SCN.

### Higher instantaneous firing frequency enhances VIP release in the SCN and circadian entrainment

Our results suggest that the firing frequency of VIP neurons mediates circadian entrainment. Higher firing frequencies (HIF vs. LIF) result in increased VIP release, larger phase shifts of the SCN and faster circadian entrainment. This optogenetic stimulation produces phase shifts consistent with the magnitude and timing of VIP-induced shifts *in vitro* (An et al., 2011) and *in vivo* (Pantazopoulos et al., 2010, Piggins et al., 1995).

Ultimately, the firing frequencies observed within VIP neurons and required for VIP-mediated phase shifting are on average 5-fold lower than frequencies that cause neuropeptide release in other neuropeptidergic neurons (Verhage et al., 1991, Iverfeldt et al., 1989). It remains to be seen whether VIP SCN neurons have a lower firing threshold to evoke VIP release or whether other factors influence VIP release. For example, daily rhythms in VIP abundance or intracellular calcium levels (Francl et al., 2010, Shinohara et al., 1999, Okamoto et al., 1991, Hastings et al., 2014) could affect release probabilities. Factors such as VPAC2 receptor abundance (An et al., 2012), and responsivity in downstream SCN neurons (Enoki et al., 2017, Fan et al., 2015) likely contribute to shaping the phase response curve and circadian entrainment following firing of VIP neurons.

### Stimulation of VIP neurons causes acute behavioral effects

Intriguingly, locomotor activity decreases during stimulation of VIP SCN neurons. This is consistent with a report correlating SCN firing with locomotor inactivity (van Oosterhout et al., 2012) and provides the first causal demonstration that firing increases within VIP SCN neurons inhibit locomotor activity. This, combined with the broad projections of patterns of VIP SCN neurons within the hypothalamus, suggests a role for VIP neurons in regulating circadian behaviors including sleep timing(Aston-Jones et al., 2001) and daily hormonal levels(Fahrenkrug et al., 2012, Loh et al., 2007). How VIP SCN neurons couple anatomically and functionally to behavioral circuits is an open question.

In conclusion, we addressed outstanding questions about how firing activity of VIP neurons alters circadian activity and behavior. By altering the instantaneous firing frequency of VIP SCN neurons, we found that firing of VIP neurons had a larger effect on circadian entrainment when stimulated with short interspike intervals, regardless of the total number of action potentials. Higher firing frequencies correlated with larger phase shifts, faster entrainment and more VIP release. We conclude that firing of VIP neurons can phase shift and ultimately entrain circadian behavior through the release of VIP.

## Methods

### Animals

VIPCre knock-in mice (VIP^tm1(cre)Zjh^, Jackson Laboratories), floxed ChR2 (Ai32, Gt(ROSA)26Sor^tm32(CAG-COP4*H134R/EYFP)Hze^, Jackson Laboratories) and *Per2::Luc(Yoo et al., 2004)* knock-in mice (founders generously provided by Dr. Joseph Takahashi, UTSW) were housed in a 12h:12h light:dark cycle in the temperature and humidity controlled Danforth Animal Facility at Washington University in St. Louis. All animals were congenic on a C57BL/6JN background. VIPChR2 mice were generated by crossing VIPCre knock-in mice to floxed ChR2 mice (Ai32). The cre within VIP neurons interacted with the loxP sites surrounding a stop codon allowing for expression of ChR2 solely within VIP neurons. Combinations of these genotypes were used for all experiments, with VIPChR2 animals being heterozygous for both VIPCre and floxed-ChR2 and controls being littermate animals heterozygous for only VIPCre or floxed-ChR2. Mice were genotyped by PCR before and by presence (ChR2-positive) or absence of eYFP fluorescence microscopy following each experiment. All procedures were approved by the Animal Care and Use Committee of Washington University and followed National Institutes of Health guidelines.

### Multielectrode array cell electrophysiology

Homozygous VIPCre-J mice were crossed with homozygous floxed-ChR2 mice to generate heterozygous litters expressing ChR2 solely in VIP+ neurons (VIPChR2). Extracellular recordings were made from multielectrode arrays (Multichannel Systems, Reutlingen, Germany) plated with SCN cells as previously described (Aton et al., 2005, Webb et al., 2012, Freeman et al., 2013). Briefly, following decapitation, the brains were rapidly removed from postnatal day 4-5 (P4-P5) VIPChR2 pups. We dissected the bilateral SCN from 250-um thick coronal brain slices, papain-dissociated and dispersed the cells at high density onto sixty, 30μm-diameter electrodes (200μm spacing) pre-treated with poly-D-lysine/laminin. Cultures were maintained in Air-DMEM (Dulbecco’s Modified Eagles Medium, DMEM, supplemented with 10% fetal bovine serum for the first week of recording) for 3 weeks prior to recording.

Multielectrode arrays were covered with a fluorinated ethylene-polypropylene membrane before transfer to a recording incubator maintained at 36°C. We waited 24h before the start of digitization to ensure culture health and stability. For extracellular recordings, spikes that exceeded a manually set threshold (~3-4 standard deviations from noise level) were digitized (1 ms before and after crossing the threshold; MC-Rack software, Multichannel Systems) at 20,000 Hz sampling. Subsequently, the culture was stimulated with 15ms pulses of light from a high-power 470nm LED (Cree XLamp XP-E2 Blue High Power LEd, LEDsupply) at 2-20Hz for 1 h with an intensity of 5 - 10 mW.

### Whole-cell patch clamp electrophysiology

Whole-cell, patch-clamp recordings from SCN neurons were obtained using procedures described previously (Hermanstyne et al., 2016). Specifically, SCN slices were prepared from 3-month old, adult, VIPChR2 heterozygous mice. After anesthesia with 1.25% Avertin, brains were removed into a cold cutting solution (in mM: 240 sucrose, 2.5 KCL, 1.25 NaH_2_Po_4_, 25 NaHCo_3_, 0.5 CaCl_2_ and 7 MgCl_2_, saturated with 95% O_2_/5%Co_2_). 300um coronal slices were cut on a Leica VT1000 S vibrating blade microtome and incubated in oxygenated artificial cerebrospinal fluid (in mM: 125 NaCl, 2.5 KCL, 1.25 NaH_2_PO_4_, 25 NaHCO_3_, 2 CaCl_2_, 1 MgCl2, 25 dextrose, saturated with 95% O2/5% CO2) for at least 1 h. Using glass pipettes (4-7 MΩ) containing an intracellular solution (in mM: 120 KMeSO_4_, 20 KCl, 10 HEPES, 0.2 EGTA, 8 NaCl 4 Mg-ATP 0.3 Tris-GTP, and 14 phosphocreatine), a “loose patch” cell-attached recording was obtained. A gigaOhm seal (>2 GΩ) was formed and spontaneous firing was recorded for approximately 1 min. We then evoked firing in ChR2-positive neurons with 15ms pulses from a 465 nm laser (DPPS MDL-III-447 100mW, 5% stability, Information Unlimited) positioned over the slice controlled by a TTL input from a Grass stimulator (S88, Grass Instrument Company, Quincy, MA) at the desired frequency (2 to 15Hz). Electrophysiological data were compiled and analyzed using ClampFit, Mini Analysis, and Prism 7.0 (Graphpad Software, La Jolla California).

### Isolation of individual neuronal firing patterns

Software was created for semiautomated spike sorting to allow discrimination of neuronal activity from MEA recordings across multiple days. Briefly, time-stamped spikes from a given electrode were separated into 24-h epochs, subsampled by taking a random 10% of the total spikes from each epoch, and then sorted based on principal component analysis (PCA) and fitting a Gaussian mixture model (GMM) to the principal components that contained >10% of the explained variance using a Bayesian information criterion (BIC) cutoff. The noise was identified as the cluster with the average spike shape with the lowest magnitude. Every non-noise cluster was then separated, and PCA and GMM clustering were applied recursively to the individual clusters to ensure coherence. This procedure (PCA, GMM clustering, separating the clusters) was repeated until each cluster could not be split based on the BIC cutoff. We used the Mahalanobis distance to keep only spikes sufficiently close to the center of the distribution. The PCA components and GMM parameters used to sort the subsampled day of spikes were saved and used to sort all spikes on that electrode from that day. The spike trains identified on each electrode were then combined across days by correlating spike shapes (Pearson r >0.95) to recover the activity of a single neuron throughout the multiday recording. If neuronal firing could not be connected across multiple days, it was excluded from subsequent analysis. Firing activity during optogenetic stimulation was similarly sorted and matched to the third day of spontaneous firing to identify VIP+ neurons. This algorithm was constructed in the Python language using packages scipy (Walt et al., 2011), and scikit-learn (Pedregosa et al., 2011) for sorting, and neuroshare for reading raw MEA data files. All scripts used in spike sorting are publicly available at: http://github.com/JohnAbel/spikesort. This method produced similar numbers of circadian neurons and spike times to manual sorting (Freeman et al., 2013) in approximately 90% less time. To calculate circadian rhythmicity, we binned the average firing rate of each neuron over 10min. Using the MetaCycle package (Wu et al., 2016) (https://cran.r-project.org/web/packages/MetaCycle/index.html in R), we calculated circadian rhythmicity using JTK cycle and Lomb-Scargle (range 20 to 28h). If a neuron was deemed rhythmic with p < 0.05 after controlling for multiple comparisons on both methods, we considered that neuron circadian.

### Classification of individual neuronal firing patterns

Neuronal firing patterns from the three days of spontaneous activity were identified in Python using scipy hierarchical clustering (Johnson, 1967) of interspike interval histogram (ISIH) with a bin size of 0.01 s and the dominant instantaneous firing rate (DIFR, the peak of the ISIH), using dynamic time warping (Salvador and Chan, 2007) as the correlation metric, and complete linkage. Varying ISIH bin size (0.005 s to 0.05 s) did not alter results significantly. The threshold for dendrogram cluster identification was set to 70% of the maximum distance between data points (default scikit-learn setting). Following clustering, DFIR was identified with a bin size of 0.0001 s to construct summary statistics with high temporal resolution. In-slice recording and whole-cell patch clamp recording data were analyzed in an identical fashion, except ISIH bin size was changed to 0.05 s for patch clamp recording due to the short 60 s recording interval.

### Recording and analysis of real-time clock gene expression

To record circadian PER2 protein expression, we crossed heterozygous VIPChR2 and *PER2::Luc* mice. Control mice were littermates lacking either VIPCre or floxed-ChR2. Adult (> P15) mice of both sexes were sacrificed with CO_2_ and 300μm coronal brain slices were sectioned. The bilateral SCN was dissected out and cultured on Millicell-CM inserts (Millipore, Billerica, MA) in pre-warmed culture medium (AirDMEM supplemented with 10mM HEPES and 100uM beetle luciferin, Promega, Madison, WI). The sealed 35-mm Petri dishes (BD Biosciences, San Jose, CA) were transferred to a light-tight incubator kept at 36°C. As described previously (Freeman et al., 2013, Aton et al., 2005), bioluminescence from the Per2-luciferase reporter was counted with a photomultiplier tube (PMT; HC135-11 MOD, Hamamatsu Corp., Shizuoka, Japan) in 6-min bins for at least three days prior to optogenetic stimulation. PMT recordings were paused during 1 h of optogenetic stimulation between CT9-12 for either one day or three consecutive days. Using a custom-made LED array (Cree XLamp XP-E2 Blue Color High powered LEDs, LEDsupply, Randolph, VT) that delivered light flashes (15 ms, 5 mW, 470 nm) per dish, SCN were stimulated for 1 h at either high (20 Hz pulses at 2 Hz) or low instantaneous frequencies (4 Hz). The LED array was powered by a supply (LEDD1B high-powered LED driver, Thorlabs, Newton, NJ) under TTL control from a stimulator (S88, Grass Instrument Company, Quincy, MA). Care was taken to minimize any mechanical disturbance to the SCN, and during stimulation, the temperature underneath the LED array remained at 36°C. After stimulation, the dishes were repositioned underneath their PMT and recording continued for at least four days.

For pharmacology experiments, we stimulated VIPChR2 PER2 slices in the presence or absence of 10uM VPAC2R antagonist ([D-p-Cl-Phe6,Leu17]-VIP, Tocris Bioscience, Bristol, UK) in AirDMEM. Following 1h of stimulation with HIF, all slices were transferred briefly into a fresh, prewarmed dish of AirDMEM and then transferred back into their original dishes and placed under their PMT channels. Application of VPAC2R antagonist alone has been previously shown to not produce phase shifts. (Jones et al., 2015)

All data were analyzed blinded to genotype. SCN traces that did not retain rhythmicity due to media evaporation or fungal infection were excluded (N=4), and all other data were analyzed in a stereotyped, reproducible manner. Each experiment (HIF stimulation, LIF stimulation and antagonist treatment) represents at least 3 separate runs with control and experimental conditions run in parallel in the same incubator. Raw counts from the PMT were detrended using a running 24-hour smooth, discarding the first and last 12 h of the recording as previously described (Herzog et al., 2015). For single-pulse experiments, the data from the day of stimulation also was excluded from analyses due to stimulation artifacts accentuated by detrending. Circadian period was calculated from a linear fit to times of the daily acrophase of PER2 expression (baseline= second through fourth days of recording; after-stimulation= fifth through seventh days; Clocklab, Actimetrics). The difference between the baseline-extrapolated and observed acrophases in the three days following stimulation was reported as the phase shift. For PMT traces stimulated for three consecutive days, we used the rising phase as a stable phase marker. Given the natural spread of PER2, we normalized the data based on the rising phase on the day before stimulation.

### Detecting VIP release after optogenetic stimulation

VIP sensor production, recording, and analysis was performed (Jones et al., 2018). Briefly, HEK293 cells were sequentially transfected with pcDNA3.1(+)−hVIPR2 and pcDNA3.1(+)-hGa15(16) (cDNA Resource Center) and selected using 0.4 μg/ml puromycin followed by 550 μg/ml G418 sulfate (Thermo Fisher Scientific). 24 h before recording, 5 × 10^4^ cells were plated on glass-bottom dishes coated with a mixture of poly-D-lysine, laminin, and collagen (Sigma). 3-4 h after plating, cells were transfected pGP-CMV-NES-jRCaMP1b (Addgene, Dana et al. 2016) and switched to recording media (DMEM supplemented with 100 U/ml penicillin, 100 μg/ml streptomycin, 10 mM HEPES, 0.56 g/L NaHCO3, 4.5g/L D-glucose, 2% L-glutamine, and 1x B27 supplement, Thermo Fisher Scientific). Organotypic SCN slices were obtained as described above, incubated overnight at 37°C, and transferred to dishes containing cultured sensor cells for recording. Recordings were performed at CTs 12-14 (extrapolated from the prior light cycle). Baseline VIP sensor fluorescence was imaged at 1 Hz for 5-8 min using a TRITC filter cube (Chroma) and a digital CCD camera (QIClick, QIImaging) with QCapture and CamStudio software. Slices were then presented with 1 min HIF or LIF stimulation (~2 mW at slice, DPPS MDL-III-447) and subsequently recorded for an additional 8-10 min. For analysis, movie frames were opened in ImageJ and bleach corrected using an exponential fit and background subtracted using a rolling ball algorithm. Regions of interest underneath the SCN were selected using a standard deviation projection and diameter thresholds of 5-30 pixels^2^. Change in fluorescence over baseline fluorescence (dF/F) for each ROI was calculated as (pixel intensity value for each frame - median pixel intensity value across all frames) / (median pixel intensity value across all frames) and smoothed using a 5-s moving average filter. Maximum dF/F for each cell was then calculated before and after stimulation.

### *In vivo* stimulation of VIP+ neurons

To stimulate VIP+ neurons *in vivo*, mice underwent stereotactic surgery to implant a fiberoptic cannula capable of delivering light to the bilateral SCN. Specifically, anesthetized mice (2% Isofluorane) were placed into a stereotaxic device and implanted with a sterilized fiber-optic cannula (5.8 mm in length, 200uM diameter core, 0.39 NA; Thorlabs, Newton, NJ). The cannula was implanted at +0.4mm anterior, +0.0 mm lateral and −5.5 mm ventral to Bregma. Mice received analgesic treatment during recovery and were enucleated as previously described (Aton et al., 2004, Hermanstyne et al., 2016) so that they did not respond to ambient light. Mice were tethered to a flexible fiberoptic cable (Thorlabs, Newton, NJ) attached to a laser (465 nm, 100 mW, 5% stability, DPPS MDL-III-447, Information Unlimited) which delivered stimuli for up to 40 days. Mice received HIF or LIF for an hour daily separated by at least four days without stimulation. Stimulated groups were randomized so that half of the mice received HIF stimulation first and the other half LIF. Mice had *ad lib* access to food, water and an open-faced wheel in a custom-built cage. Wheel revolutions were counted with a reed switch (Clocklab, Actimetrics).

### Immunohistochemistry

To test for neuronal activation, we measured cFOS protein induction in mice implanted with a fiberoptic aimed at the SCN after 1 h of stimulation (15Hz, 15ms pulses) during early subjective night (CT 13). Immediately following stimulation, mice were anesthetized with 1.25% Avertin (2,2,2-tribromoethanol and tert-amyl alcohol in 0.9% NaCl; 0.025 ml/g body weight) and transcardially perfused with phosphate-buffered saline (PBS) and 4% paraformaldehyde (PFA). The brain was rapidly dissected and transferred to 30% sucrose following 24 h in 4% PFA. Frozen coronal sections cut at 40 um were collected in 3 separate wells. For cFOS immunofluorescence, free-floating sections were washed for 1 h in PBS, incubated overnight at 4°C in rabbit anti-cFOS antibody (1:1000 in PBSGT; Santa Cruz Biotechnology, Santa Cruz, CA). Slices were washed again and incubated for 2 h at room temperature in donkey anti-rabbit Cy3 secondary antibody (1:500 in PBSGT). Sections were washed again in PBS for 30min, mounted, and cover-slipped with DABCO (1,4-Diazobicyclo[2,2,2]-octane) mounting medium. Sections were imaged in 4-um z-stacks on a Nikon A1 Confocal microscope. Two independent investigators quantified the fraction of ChR2-eYFP positive neurons that also expressed nuclear cFOS. Results differed by less than 10% per brain. For avidin-biotin immunohistochemistry, free-floating sections were incubated for 72 h in the same rabbit c-FOS antibody (1:2500). Tissues were reacted in diaminobenzidine with 0.01% H_2_O_2_, mounted, dehydrated and cover-slipped. Sections were imaged using the Alafi Nanozoomer at Washington University in St. Louis Medical School. Tissues were always processed together and mid-SCN sections were selected from all animals for quantification. An investigator blinded to the genotype of the mouse quantified the number of cFOS positive cells within the SCN using ImageJ software. The SCN was located and boundaries were drawn to demarcate the ventral and dorsal SCN in each animal (250 um ×150 um, ventral SCN per side, 150 um × 150 um, dorsal SCN per side).

### Locomotor analysis

We identified the daily onset of locomotor activity from wheel running data in 15-min bins as the zero-crossing of the continuous wavelet transform using the Mexican Hat wavelet as described for temperature rhythms (Leise et al., 2013). We calculated the phase response curve by identifying the change in activity onsets evoked by optogenetic stimulation compared to the same mouse under free running conditions. Calculating phase change in comparison to the same mouse under free-running conditions (rather than comparison with a separate control group) allowed inclusion of mice with visibly different periods. To measure the time to entrainment, we constructed Rayleigh plots of activity onset for days 2-10 of optogenetic stimulation and calculated the Kuramoto parameter (i.e. synchronization index)(Kuramoto, 2012). Finally, we measured optogenetically induced activity suppression by comparing wheel revolutions during the stimulated hours between CT 12 – 16 and equivalent time bins from free-running days with no stimulation.

## Author Contributions

C.M. and E.D.H. designed the research.

C.M. performed the MEA, PMT *in vitro*, and locomotor *in vivo* experiments.

C.M. and S.C. performed the *in vivo* cFOS experiments.

T.H. performed the whole-cell optogenetics experiments.

J.H.A. and C.M designed and wrote the spike sorting algorithm with guidance from F.J.D. and E.D.H.

J.H.A., C.M., and E.D.H. analyzed the MEA, in-slice, and whole cell electrical activity data.

C.M. and S.C. analyzed the cFOS data.

C.M., J.H.A., and E.D.H. analyzed the *in vivo* locomotor data.

T.H. and E.D.H. analyzed the whole cell optogenetics data.

C.M., J.H.A., and E.D.H. wrote the paper.

J.J. and T.S. performed the VIP Sniffer experiments.

J.J. analyzed the VIP Sniffer experiments.

## Acknowledgements

This work was supported by NIH grants NS095367 (EDH), F31-GM11517 (CM), T32- HLO7901 (JHA), and the Hope Center for Neurological Disorders at Washington University. The authors thank Dr. Hanspeter Herzel, Dr. Bharath Ananthasubramaniam, and Kelsey R. Dean for valuable discussions.

